# A survey of public attitudes towards third-party reproduction in Japan in 2014

**DOI:** 10.1101/328229

**Authors:** Naoko Yamamoto, Tetsuya Hirata, Gentaro Izumi, Akari Nakazawa, Shinya Fukuda, Kazuaki Neriishi, Tomoko Arakawa, Masashi Takamura, Miyuki Harada, Yasushi Hirota, Kaori Koga, Osamu Wada-Hiraike, Tomoyuki Fujii, Minoru Irahara, Yutaka Osuga

## Abstract

*Objective:* The objective of this study was to examine public attitudes towards third-party reproduction and the disclosure of conception through third-party reproduction.

*Methods:* We conducted the web-based survey for the public attitude towards third-party reproduction in February 2014. Twenty-five hundred people were recruited with equal segregation of age (20s, 30s, 40s, and 50s) and gender. We analyzed the association between gender, age, infertility, and ethical view using a questionnaire regarding donor sperm, donor oocyte, donor embryo, gestational surrogacy, and disclosure to offspring.

*Results:* Of the respondents, 36.2% approved and 26.6% disapproved with gamete or embryo donation. The frequency of those who approved was the lowest in females in the 50-59 years age group, and was significantly higher in males or females with infertility. Secondly, 40.9% approved and 21.8% disapproved with gestational surrogacy. The frequency of those who approved gestational surrogacy was higher in males or females with infertility. Thirdly, 46.3% of respondents agreed and 20.4% disagreed with “offspring have the right to know their origin”. Those who disagreed were primarily in the 50-59 age group of both genders, and disagreement was significantly higher in the infertility group compared with non-infertility group.

*Conclusion:* In this study, public attitudes were affected by gender, age, and experience of infertility. These study findings are important in understanding the attitude towards third-party reproduction and disclosure to the offspring. Respondents having indecisive attitudes were >30%, which might indicate an increased requirement for information and education to enhance the discussion on the ethical consensus on third-party reproduction in Japan.

## Introduction

The treatment cycle of assisted reproductive technology (ART) has dramatically increased; more than 1 million babies were born by ART between 2008 and 2010 [1]. In Japan, 424151 treatment cycles were carried out in 2015, and 51001 neonates (1 in 19.7 neonates born in Japan) were born [2]. Although ART is now widely accepted as clinically effective for treatment of many forms of subfertility, some people cannot conceive due to poor reproductive function, for example, uterine infertility and poor ovarian function. For those patients, ART using third party sperm, oocytes, or uterus is an option, known as third-party reproduction.

It is not technically difficult to perform third-party reproduction. Indeed, in the United States, there has been many cases of third-party reproduction, accounting for 16% of the total ART [3]. However, ethical concerns have been raised about these fertility treatments. The attitudes toward third-party reproduction are different in each country, which also depends on a country’s law on whether oocyte donation is legal or not. For example, in the EU, oocyte donation is permitted under certain conditions in France, but it is forbidden in Germany and Italy. In Japan, there is no legislation concerning ART, gamete or embryo donation, and gestational surrogacy, which have been controlled by the Japan Society of Obstetrics and Gynecology guidelines [4]. As for surrogate pregnancies, the academic society prohibits academic members from being involved in surrogate pregnancies at the present time, and sperm or egg donation are not prohibited and limitedly performed.

The ethical consequences of using donated gametes, embryos, and gestational surrogacy are a matter for debate among professionals and society in general. In Japan, there have been several surveys on general attitudes toward third-party reproduction in 1999, 2003, and 2007 [5–7]. In addition, the ethical views of the general population regarding third-party reproduction are worthy of close examination. Furthermore, the matter of disclosure of donor conception to donor offspring is a very contentious issue [8]. Therefore, we sought to conduct a survey on public attitudes to third-party reproduction and the disclosure of donor conception to offspring. In this study, the association between gender, age, infertility, and the ethical view were examined in a large representative sample thorough a web-based questionnaire regarding donor sperm, donor oocytes, donor embryos, gestational surrogacy, and disclosure of donor conception.

## Materials and Methods

We conducted a web-based survey to assess the public attitude towards third-party reproduction. The sampling frame was developed by an internet research company and we sent out questionnaires regarding third-party reproduction through a website. Respondents were asked to read a summary page explaining the purpose and content of the questionnaire prior to starting the survey. In answering the questionnaire, reference materials (Supplemental Figure 1) were also included to deepen the understanding of respondents’ opinion on third-party reproduction. The questionnaire also included items regarding age, gender, marriage status, number of children, annual family income, and educational background. The web-based questionnaire was available online from 14^th^ February to 25^th^ February 2014. The criterion for exclusion from the survey was non-completion of the whole interview. Twenty-five hundred people were recruited with an equal segregation of age (20s, 30s, 40s, and 50s) and sex. The study was conducted with the approval of the ethics committee of the University of Tokyo Hospital.

### Statistical analysis

The statistical analyses were performed using JMP pro version 13 (SAS Institute Inc, Cary, NC, USA). The categorical data were analyzed using chi-square test and Fisher’s exact test, and is presented as numbers and percentages. A P value < 0.05 was considered statistically significant.

## Results

### Characteristics of respondents

The study consisted of a total of 2500 respondents with an equal number across each age group (20s, 30s, 40s, and 50s). Answers from the same number of men and women in each age group were obtained. Of the respondents, 44.3% were unmarried and 55.7% were married. Survey respondents had a wide range of family income levels. The characteristics of the respondents are shown in Table 1.

**Table 1.** Demographic data of respondents (n=2500)

|  | Total | Male | Female |
| --- | --- | --- | --- |
| Age (years) |  |  |  |
| 20-29 | 624 (25.0%) | 312 (25.0%) | 312 (25.0%) |
| 30-39 | 626 (25.0%) | 313 (25.0%) | 313 (25.0%) |
| 40-49 | 626 (25.0%) | 313 (25.0%) | 313 (25.0%) |
| 50-59 | 624 (25.0%) | 312 (25.0%) | 312 (25.0%) |
| Marital status |  |  |  |
| unmarried | 1108 (44.3%) | 649 (51.9%) | 459(36.7%) |
| married | 1392 (55.7%) | 601(48.1%) | 791(63.3%) |
| Number of children |  |  |  |
| 0 | 1343 (53.7%) | 740 (59.2%) | 603 (48.2%) |
| 1 | 407 (16.3%) | 170 (13.6%) | 237 (19.0%) |
| 2 | 561 (22.4%) | 249 (19.9%) | 312 (25.0%) |
| 3 | 164 (6.6%) | 79 (6.3%) | 85 (6.8%) |
| 4 | 21 (0.8%) | 11 (0.9%) | 10 (0.8%) |
| more than 4 | 4 (0.2%) | 1 (0.1%) | 3 (0.2%) |
| Annual family income (yen/year) |  |  |  |
| less than 1,000,000 | 145 (5.8%) | 81(6.5%) | 64 (5.1%) |
| 1,000,000-3,000,000 | 412 (16.5%) | 185 (14.8%) | 227 (18.2%) |
| 3,000,000-5,000,000 | 686 (27.4%) | 345 (27.6%) | 341 (27.3%) |
| 5,000,000-7,000,000 | 571 (22.8%) | 276 (22.1%) | 295 (23.6%) |
| 7,000,000-10,000,000 | 417 (16.7%) | 223 (17.8%) | 194 (15.5%) |
| 10,000,000-15,000,000 | 200 (8.0%) | 109 (8.7%) | 91 (7.3%) |
| 15,000,000- | 69 (2.8%) | 31 (2.5%) | 38 (3.0%) |
| Education |  |  |  |
| Junior high school | 68 (2.7%) | 36 (2.9%) | 32 (2.6%) |
| High school | 645 (25.8%) | 308 (24.6%) | 337 (27.0%) |
| Technical college | 367 (14.7%) | 177 (14.2%) | 190 (15.2%) |
| Junior college | 275 (11.0%) | 16 (1.3%) | 259 (20.7%) |
| University or graduate school | 1144 (45.8%) | 713 (57.0%) | 431 (34.5%) |
| others | 1 (0.0%) | 0 (0.0%) | 1 (0.1%) |
| Experience of infertility |  |  |  |
| yes | 358 (14.3%) | 122 (9.8%) | 236 (18.9%) |
| none | 2142 (85.7%) | 1128 (90.2%) | 1014 (81.1%) |
| Experience of infertility treatment |  |  |  |
| none | 2259 (90.4%) | 1166 (93.3%) | 1093 (87.4%) |
| timed intercourse | 206 (8.2%) | 67 (5.4%) | 139 (11.1%) |
| IUI (intrauterine insemination) | 77 (3.1%) | 24 (1.9%) | 53 (4.2%) |
| IVF-ET (In vitro fertilization - embryo transfer) | 44 (1.8%) | 17 (1.4%) | 27 (2.2%) |
| ICSI (intracytoplasmic sperm injection) | 31 (1.2%) | 13 (1.0%) | 18 (1.4%) |

### Social acceptance

Regarding social and ethical acceptance of gamete or embryo donation, the rate was 36.2% for “should be approved”, 26.6% for “should *not* be approved”, and 37.3% for “indecisive” (Table 2). The frequency of respondents who gave “should be approved” in males and females in the 20-29, 30-39, and 40-49 age groups were significantly higher than females in the 50-59 age group. The frequency of respondents who gave “should *not* be approved” in males and females in the 50-59 age group was significantly higher than in the other age groups. As shown in Table 3, the frequency of a “positive response” was higher in males or females with infertility than those without infertility. Furthermore, the frequency of respondents who were indecisive was smaller in those with infertility than those without infertility. There were no effects of marital status, number of children, or annual family income.

**Table 2.**
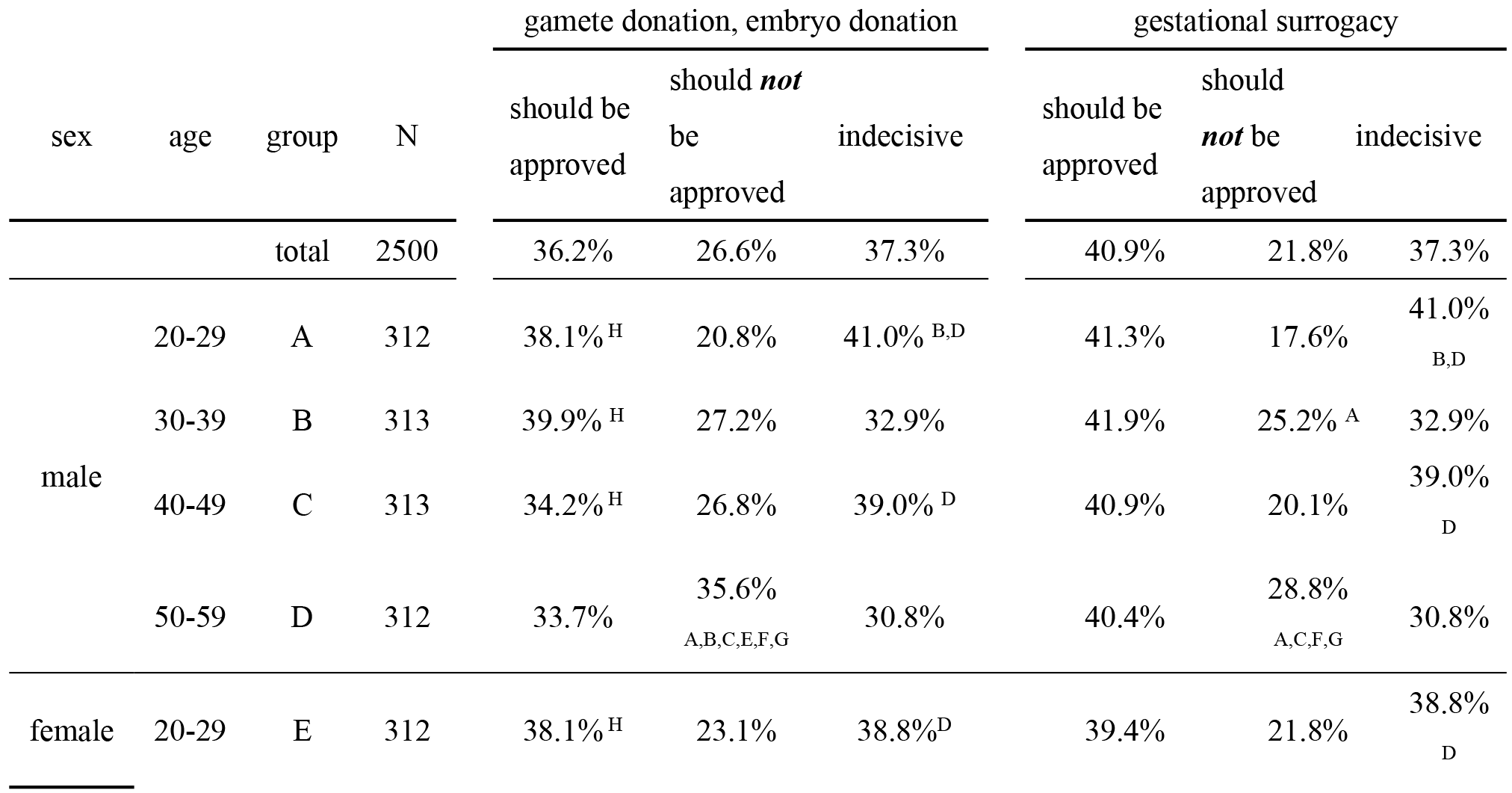
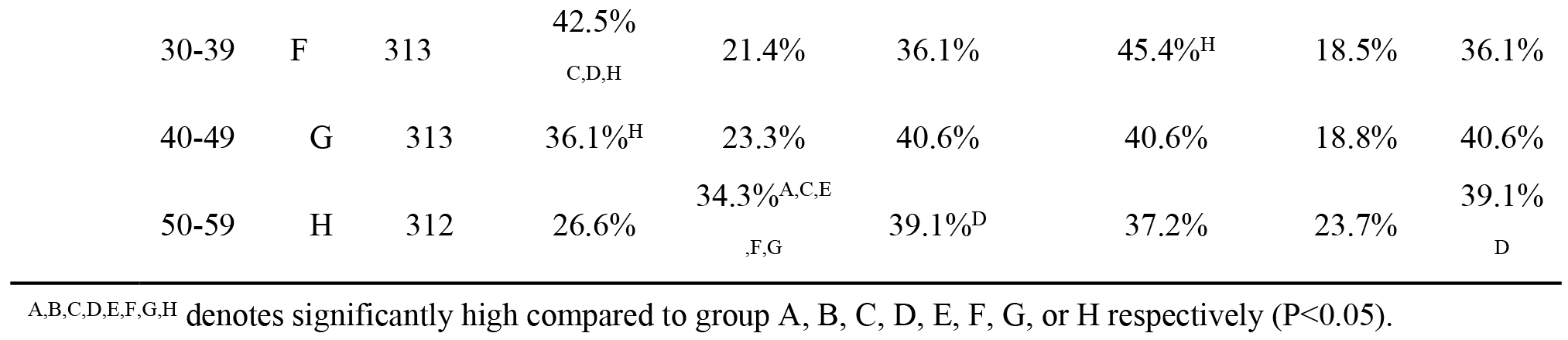
Comparison of public attitude towards gamete or embryo donation and gestational surrogacy divided by age and gender

**Table 3.** Comparison of public attitudes towards gamete or embryo donation and gestational surrogacy divided by fertility

|  |  |  |  | gametes donation, embryo |  |  |  |  |  |
| --- | --- | --- | --- | --- | --- | --- | --- | --- | --- |
|  |  |  |  | donation |  |  | gestational surrogacy |  |  |
| sex | infertility | group | N | should<br>be<br>approved | should<br>*not* be<br>approved | indecisive | should<br>be<br>approved | should<br>*not* be<br>approved | indecisive |
| total |  |  | 2500 | 36.2% | 26.6% | 37.3% | 40.9% | 21.8% | 37.3% |
| male | + | A | 122 | 45.1%<br>B,D | 32.8% | 22.1% | 50.8%<br>B,D | 27.0% | 22.1% |
|  | - | B | 1128 | 35.5% | 27.0% | 37.4%<sup>A,C</sup> | 40.1% | 22.5% | 37.4%<sup>A,C</sup> |
| female | + | C | 236 | 44.9%<br>B,D | 27.1% | 28.0% | 51.7%<br>B,D | 20.3% | 28.0% |
|  | - | D | 1014 | 33.7% | 25.1% | 41.1%<sup>A,C</sup> | 38.1% | 20.8% | 41.1%<sup>A,C</sup> |

<sup>A,B,C,D</sup> denotes significantly high compared to group A, B, C, or D respectively (P<0.05).

As for surrogacy, 40.9% of respondents agreed that it “should be approved”, 21.8% responded with “should *not* be approved”, and 37.3% were “indecisive” (Table 2). The frequency of those who approved gestational surrogacy was around 40% in any age and gender group. The frequency of those who disapproved was highest in males in the 50-59 age group. As shown in Table 3, the frequency of those who approved gestational surrogacy was higher in males or females with infertility than in those without infertility, which was similar to the attitudes towards gamete or embryo donation.

Next, we asked the question “who is eligible to be a recipient of gametes or embryos?”. In response, 88.6%, 33.6%, 13.4%, 10.0%, and 25.8% answered that a couple, a fact-married couple, a single woman, a single man, or a homosexual couple are acceptable as recipients, respectively (Table 4). The frequency of those who answered it as acceptable for a homosexual couple was higher in females in the 20-29, 30-39, and 40-49 age groups than in males and females in the 50-59 age group. There were no effects of marital status, number of children, annual family income, or infertility.

**Table 4.**
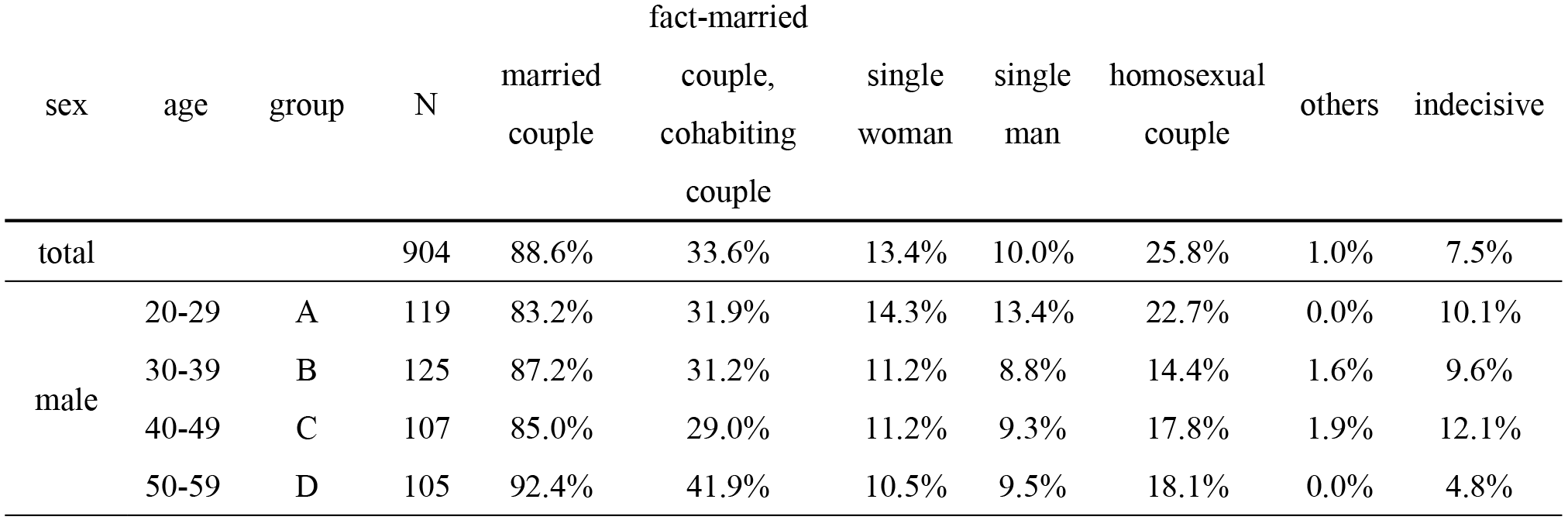
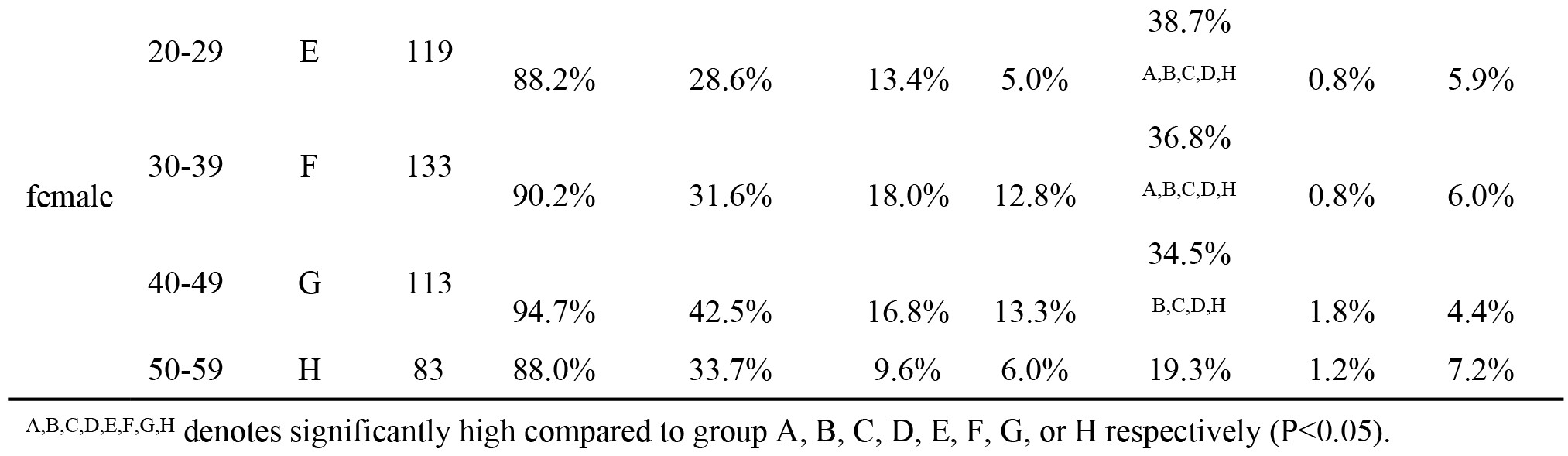
Who is eligible to be a gamete or embryo recipient?

The most frequent reasons why respondents thought that gamete or embryo donation was socially acceptable were: "because there is a possibility that a person who cannot conceive due to disease can have a child" (79.6%), and "because people who are unable to conceive because of aging will be able to be pregnant"(43.9%) (Supplemental Table 1). In addition, the most frequent reasons why respondents thought that gamete or embryo donation was *not* socially acceptable were: “because there is no genetic link with parenting parents” (46.8%), and “parent-child relationship will become unnatural” (39.9%) (Supplemental Table 2). As for surrogacy, the reasons why respondents thought that gestational surrogacy was socially acceptable were: “because there is a possibility that a person who cannot conceive due to illness can have a child” (76.7%), and “because there is a possibility that a woman who removed the uterus due to disease or accident can have a child” (66.6%) (Supplemental Table 3). Furthermore, the reasons why respondents thought that gestational surrogacy was *not* socially acceptable were “parent-child relationship will become unnatural” (33.9%), and “pregnancy should be natural” (32.8%) (Supplemental Table 4).

### Personal opinion

To obtain a personal opinion, we asked whether respondents would choose to receive each type of third-party reproduction, assuming that they could not conceive in all other ways. For third-party reproduction, 1-3% would use it across all age groups; no significant difference was observed between each group. The frequency of those who would want to receive donor sperm, egg, embryo, or gestational surrogacy if their spouse wishes to was significantly higher in males than in females (P < 0.01). The frequency of those who did not want to use third party reproduction was significantly lower in males than in females (P<0.01). The rate of those who would want to use donor sperm if their spouse wishes to was 33.3%, 30.7%, 23.6%, and 22.4% in males in the 20-29, 30-39, 40-49, and 50-59 age groups, respectively, and 26.6%, 21.4%, 15.3%, and 11.9% in females in the 20-29, 30-39, 40-49, and 50-59 age groups, respectively, showing a decreasing tendency with age. The frequency of those who would want to use third party reproduction if their spouse wishes to was significantly lower in females in the 50-59 age group than in any other group. On the other hand, the rate of those who did not want to use donor sperm was 64.4%, 66.8%, 75.4%, and 76.3% in males in the 20-29, 30-39, 40-49, and 50-59 age groups, respectively, and 72.1%, 77.0%, 82.4%, and 86.5% in females in the 20-29, 30-39, 40-49, and 50-59 age groups, respectively, showing an increasing tendency with age. These trends were common in the donor egg, donor embryo, and gestational surrogacy categories, as shown in Table 5. On the other hand, there was no significant difference in attitude towards gamete or embryo donation and surrogate pregnancy between the respondents with or without infertility of either gender (Table 6).

**Table 5.**
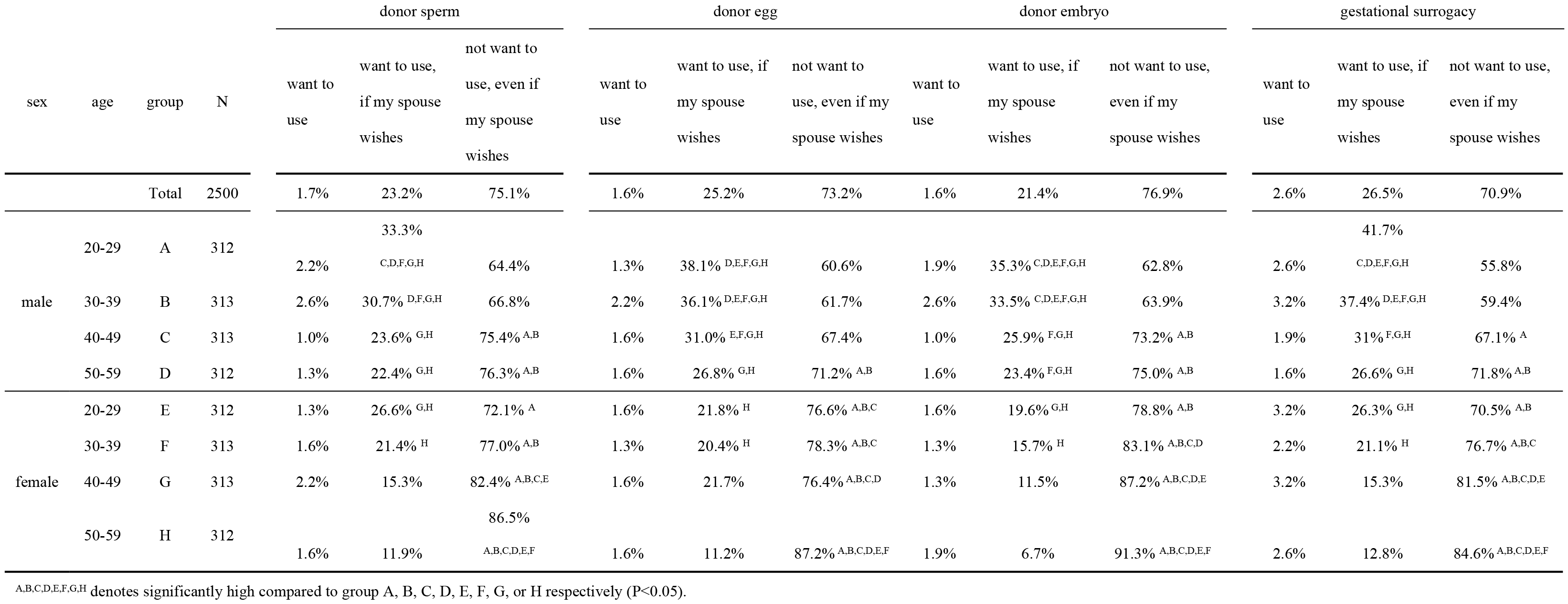
Attitudes towards each type of third-party reproduction assuming the respondents cannot conceive in other ways

**Table 6.**
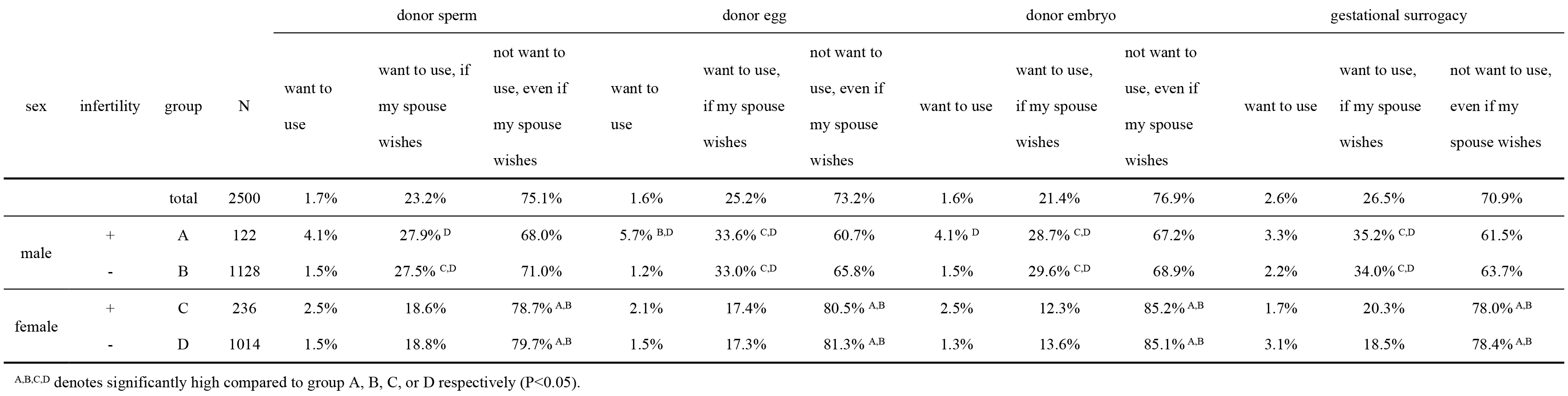
Attitudes towards each type of third-party reproduction assuming the respondents cannot conceive in other ways

### Offspring’s right to know their origin

First, we asked the respondents if we should disclose the offspring’s origin to those born via third-party reproduction; 46% of all respondents agreed, 20% disagreed, and 31% were indecisive (Table 7). The frequency of those who agreed was significantly higher in males or females in the 20-29 and 30-39 age groups than in females in the 50-59 age group, and was highest in females in the 20-29 age group (55.4%). The frequency of those who disagreed was higher in males or females in the 50-59 age group than in the 20-29 and 30-39 age groups. In addition, the frequency of those who disagreed was higher in males or females with infertility than in those without infertility (Table 8).

**Table 7.** Attitudes towards offspring’s right to know their origin

| sex | age | group | N | Offspring has the right to know their origin |  |  |  | Do you receive treatment, even if offspring are granted the right to know their origin. |  |  |  |
| --- | --- | --- | --- | --- | --- | --- | --- | --- | --- | --- | --- |
|  |  |  |  | agree | disagree | indecisive | others | YES | hesitate or<br>cease | would not<br>receive it<br>originally. | indecisive |
|  |  | total | 2500 | 46.3% | 20.4% | 31.7% | 1.6% | 18.4% | 9.5% | 39.2% | 32.9% |
| male | 20-29 | A | 312 | 48.7%<sup>H</sup> | 14.7% | 34.6% | 1.9% | 21.5%<sup>H</sup> | 10.9% | 29.8% | 37.8%<sup>D,F,G,H</sup> |
|  | 30-39 | B | 313 | 49.8%<sup>H</sup> | 18.5% | 31.0% | 0.6% | 21.7%<sup>H</sup> | 9.6% | 29.7% | 39%<sup>D,F,G,H</sup> |
|  | 40-49 | C | 313 | 42.2% | 23%<sup>A,E</sup> | 33.5% | 1.3% | 15.7% | 8.9% | 37.4% | 38%<sup>D,F,G,H</sup> |
|  | 50-59 | D | 312 | 43.3% | 27.2%<sup>A,B,E,F,G</sup> | 28.5% | 1.0% | 19.9%<sup>H</sup> | 11.2% | 39.4%<sup>A,B</sup> | 29.5% |
| female | 20-29 | E | 312 | 55.4%<sup>C,D,F,G,H</sup> | 13.1% | 29.5% | 1.9% | 21.8%<sup>G,H</sup> | 9.6% | 33.7% | 34.9%<sup>H</sup> |
|  | 30-39 | F | 313 | 47.3%<sup>H</sup> | 18.5% | 32.9% | 1.3% | 18.8%<sup>H</sup> | 9.9% | 42.2%<sup>A,B,E</sup> | 29.1% |
|  | 40-49 | G | 313 | 45.7% | 18.5% | 32.9% | 1.3%<sup>B</sup> | 15.3% | 6.7% | 47.9%<sup>A,B,C,D,E</sup> | 30.0% |
|  | 50-59 | H | 312 | 37.8% | 29.5%<sup>A,B,E,F,G</sup> | 31.1% | 1.6% | 12.2% | 9.3% | 53.8%<sup>A,B,C,D,E,F</sup> | 24.7% |

<sup>A,B,C,D,E,F,G,H</sup> denotes significantly high compared to group A, B, C, D, E, F, G, or H respectively (P<0.05).

**Table 8.** Attitudes towards offspring’s right to know their origin

| sex | infertility | group | N | Offspring has the right to know their origin |  |  |  | Do you receive treatment, even if offspring are granted the right to know their origin? |  |  |  |
| --- | --- | --- | --- | --- | --- | --- | --- | --- | --- | --- | --- |
|  |  |  |  | agree | disagree | indecisive | others | YES | hesitate or cease | would not receive it originally. | indecisive |
|  |  | total | 2500 | 46.3% | 20.4% | 31.7% | 1.6% | 18.4% | 9.5% | 39.2% | 32.9% |
| male | + | A | 122 | 45.9% | 28.7%<br>B,D | 25.4% | 0.0% | 27%<sup>B,D</sup> | 14.8%<sup>D</sup> | 35.2% | 23.0% |
|  | - | B | 1128 | 46.0% | 20.0% | 32.6%<sup>C</sup> | 1.3% | 18.9%<sup>D</sup> | 9.7% | 34.0% | 37.5%<sup>A,C,D</sup> |
| female | + | C | 236 | 49.2% | 29.7%<sup>B,D</sup> | 19.9% | 1.3% | 26.7%<sup>B,D</sup> | 12.7%<sup>D</sup> | 41.5%<sup>B</sup> | 19.1% |
|  | - | D | 1014 | 46.0% | 17.7% | 34.1%<sup>C</sup> | 2.3% | 14.8% | 8.0% | 45.1%<sup>A,B</sup> | 32.1%<sup>A,C</sup> |

<sup>A,B,C,D</sup> denotes significantly high compared to group A,B,C, or D respectively (P<0.05).

Next, we asked all respondents the question: “would you receive the third-party treatment if origin was disclosed to the offspring, assuming that you would undergo third-party reproduction?”. In response, 18.4% said yes, 9.5% would hesitate or cease to receive the treatment, 39.2% would not receive the treatment originally, and 32.9% were indecisive (Table 7). The frequency of those who would receive treatment was significantly lower in females in the 50-59 age group than in males or females in the 20-29, 30-39 age group. The rate of those who would not receive the treatment originally was 33.7%, 42.2%, 47.9%, and 53.8% in females in the 20-29, 30-39, 40-49, and 50-59 age groups, respectively, showing an increasing tendency with age.

As shown in Table 8, the frequency of those who would receive treatment was significantly higher in males or females with infertility than in those without infertility. In addition, the frequency of those who would hesitate or cease to receive treatment if the offspring were to be informed of the origin was also significantly higher in females with infertility than in those without infertility. There was no significant difference between males with or without infertility, although similar trends were observed (14.8% vs. 9.7%). The rate of those who were indecisive was significantly lower in males or females with infertility than in those without.

## Discussion

We conducted a web-based survey on the public attitude towards third-party reproduction and the disclosure of third-reproduction to the offspring in Japan. To date, there has been no large-scale survey on the public attitude towards disclosure of third-reproduction to the offspring in Japan. Firstly, approximately 40% of respondents agreed that gamete donation or gestational surrogacy should be approved, whereas 25% answered that it should not be approved. The rate of respondents who agreed was the lowest in females in the 50-59 age group, and the rate of those who disagreed was the highest in females in the 50-59 age group. In addition, the frequency of those who agreed was significantly higher in males or females with infertility, compared to those without. Secondly, the rate of those who would receive third-party reproduction if their spouse wishes to showed a decreasing trend with age, and those against receiving third-party reproduction, even if their spouse wishes to, showed an increasing trend with age. In addition, there was no significant difference in the attitudes toward gamete or embryo donation and surrogate pregnancy between the respondents with or without infertility of both genders. Thirdly, 46.3% of respondents agreed that offspring have the right to know their origin, whereas 20.4% disagreed. Those who disagreed was highest in the 50-59 age group of both genders. In addition, those who disagreed was higher in the infertility group compared with non-infertility group. In this study, we examined in detail whether gender, age, and infertility affected attitudes towards third-party reproduction. The findings of this study are important in understanding the attitudes towards third-reproduction and the disclosure to offspring.

Approximately 40% of respondents agreed that gamete or embryo donation and surrogate pregnancy should be accepted socially, which was equivalent to the 2003 survey. Although 10 years have passed, there seems to be little change in the distribution of attitudes. Furthermore, the Japan Society of Obstetrics and Gynecology guidelines prohibit surrogacy [4, 9], but over 40% of people approve gestational surrogacy.

Analysis by age revealed that the number of people who agree gamete or embryo donation has decreased in elderly people; disapproval was highest in the 50-59 age group. With regards to surrogate pregnancy, disapproval was significantly higher in men that were older in age; however, there was no significant difference depending on age among females. Respondents who had experience of infertility often answered that both gamete or embryo donation and surrogate pregnancy should be accepted, suggesting that infertility experience has an effect on individual opinion.

The top reason for agreeing with donation was that people who cannot conceive due to disease or aging may become pregnant. The main reasons for not agreeing with donation were that there was no genetic linkage and that family relations became unnatural. Oocyte donation pregnancies have been reported to be associated with adverse obstetric and neonatal outcomes [10, 11]; however, these issues were not the main reason for disapproval in this study.

Regarding gestational surrogacy, the top reason for approval was that people who cannot conceive due to disease or those who underwent a hysterectomy due to disease or accident can have children. On the other hand, the top reasons for not approving were that familial relations are unnatural and that pregnancy should be natural. Gestational surrogacy has been reported to increase adverse perinatal outcomes, including preterm birth, low birth weight, hypertension, gestational diabetes, and placenta previa, compared with spontaneous pregnancy by the same women [12]. Although this fact can be a reason for not approving of gestational surrogacy, it was not main reason for disapproval in this study. In order to conduct a survey on surrogacy, information on these risks of gestational surrogacy should be given. Indeed, a previous report also suggested that the disapproval rate of gestational surrogacy was increased by recognizing the perinatal adverse outcomes of surrogacy [7].

We also asked if the respondents would choose third-party reproduction if they could not get pregnant in other ways, which resulted in the positive opinion frequency for third-party reproduction being higher in males. This is in line with previous reports that the rate of a positive people of third-party reproduction was higher in men [6, 13, 14], which was presumed to be due to the fact that men are not involved physically in the procedure and that the desire for descendants is strong [14]. As age increased, the attitudes towards these treatments became negative, which was common to all third-party reproductions. These trends were consistent with previous reports [5]. Meanwhile, there was no significant difference in the attitude towards third-party reproduction with or without infertility, suggesting that the experience of infertility has little effect on personal choice. People with infertility tended to socially approve third-party reproduction, but when it came to themselves, they were very cautious of whether to choose third-reproduction or not. Regarding if we should disclose their origin to the offspring born with third-party reproduction, as many as 46% of respondents approved, 20.4% disapproved, and 31.7% were undecided. This prevalence of opinions is similar to the result (38.4%, 29.6%, and 32%, respectively) obtained in Germany [15], where egg donation and reception are prohibited by law. In Sweden, 83% of women and 75% of men supported the right of the offspring to know the origins [16]. This high proportion of approval might relate to the legal liberalization of gamete donation, considering that Sweden is one of the countries permitting oocyte donation and granting offspring the right to know the identity of the donor.

When analyzing the results by age and gender, the frequency of those who approved was the lowest in women in their 50s; about one-third of male or female respondents in their 50s opposed it. In addition, regarding men and women with infertility, there was a significantly higher negative opinion on the offspring’s right to know their origin, compared to those without. Furthermore, when the right to know the origin was recognized, there was a significant increase in both those who would continue treatment and those who would hesitate in those with infertility. As the number of people who answered "cannot decide" significantly reduced in those with infertility, it suggests that people with infertility can form a clear opinion whether they are positive or negative. However, over 30% were indecisive, which might indicate that it is generally difficult to visualize the consequences of different scenarios for recipients and their offspring. These findings may indicate the need for more adequate information and education of the community to enhance the discussion on the ethical consensus on third-party reproduction. Recently, the number of Japanese infertile patients who have travelled across the border to undergo third party reproduction has increased, although it is not a mainstream issue [17]. This is because third-party reproduction is not readily available to infertile Japanese patients. Several papers point out that a law or regulation on third-party reproduction should be made [4, 9]. Semba et al. indicated that Japan will merely export ethical issues to a permissive country if the banning of surrogacy in Japan results in more infertile patients going abroad for third-party reproduction [4].

Very recently, there have been several cases of birth after a uterus transplantation [18–20]. Uterine transplantation has not been carried out in Japan; however, a positive result has been obtained in a public survey on uterus transplantation in Japan, compared to gestational surrogacy [21]. Although uterine transplantation is still in an experimental phase, it can be an option and may have the potential to change the ethical viewpoints and attitudes towards gestational surrogacy.

The present study has certain limitations. First, this study is a cross-sectional study, which cannot explore a causal relationship. Second, there is the possibility of survey selection bias and issues of generalizability. After reading the questionnaire introduction, respondents had the choice whether to complete the survey or not. Third, we have not confirmed the level of knowledge and understanding on third-party reproduction of the respondents. Despite these limitations, by conducting a web-based questionnaire on the general population, we collected a large sample size; therefore, the findings of this study reflect the public attitudes towards third-party reproduction in Japan.

## Conclusion

In the current study, we clarified that the public attitudes towards third-party reproduction and the disclosure to offspring were affected by gender, age, and the experience of infertility. Respondents having indecisive attitudes were over 30%, which might indicate the requirement for more adequate information and education of the community to enhance the discussion on the ethical consensus on third-party reproduction. Investigating and gaining knowledge regarding the general attitudes towards these issues is important. Legal and medical professionals require this kind of research in order to make predictions about behaviors in the future and to make a regulation or legislation concerning third-party reproduction.

## Acknowledgements

The authors would like to thank Editage (www.editage.jp) for the English language review.

## Supporting Information Captions

S1 Figure. reference materials to deepen the understanding of respondents.

S1 Table. Why do you think that we should approve in vitro fertilization with the provision of a third-party egg, sperm, or embryo?

S2 Table. Why do you think that in vitro fertilization using sperm, egg, embryo of a third party is not socially acceptable?

S3 Table. Why do you think that we should approve a surrogate pregnancy using the womb of a third party?

S4 Table. Why do you think that surrogate pregnancy using a third party’s uterus is not socially acceptable?

